# Brain dynamics of attentional, default-mode and limbic networks are disrupted at rest in Post-COVID-19 Syndrome

**DOI:** 10.64898/2026.01.12.699034

**Authors:** Marie-Stephanie Cahart, Owen O’ Daly, Ziyuan Cai, Nicole Mariani, Alessandra Borsinni, Valeria Mondelli, Brandi Eiff, Silvia Rota, Timothy Nicholson, Laila Raida, Adam Hampshire, Ottavia Dipasquale, Lia Fernandes, Federico Turkheimer, Steven C.R. Williams, Daniel Martins

## Abstract

**Background:** Post-COVID-19 Syndrome (PCS) is characterised by persistent fatigue, cognitive impairments, and affective symptoms, yet its underlying neural mechanisms remain poorly understood. While static neuroimaging studies have identified resting-state connectivity abnormalities in PCS, such approaches fail to capture the brain’s dynamic functional organisation. This represents a missed opportunity to understand how alterations at the dynamic macroscale interactions give rise to the complex and fluctuating symptom profile of PCS. Cognitive and emotional processes rely on the brain’s capacity to flexibly reconfigure large-scale networks over time; disruptions in the exploration and transition between brain states may therefore play a central role in PCS pathophysiology.

**Methods:** Resting-state fMRI data were acquired from 20 individuals with PCS (mean age = 41.8 years, SD = 9.4) and 20 age- and sex-matched healthy controls (mean age = 40.6 years, SD = 8.1) using a multi-echo sequence. Following denoising with multi-echo independent component analysis, we applied Leading Eigenvector Dynamics Analysis (LEiDA) to identify recurrent patterns of whole-brain phase synchrony. The optimal number of dynamic brain states was determined using the Dunn index. For each state, we quantified probability of occurrence, lifetime, and transition probabilities, and mapped spatial topographies onto canonical functional networks. Group differences were assessed using ANCOVAs controlling for age, sex, and handedness. Exploratory associations with clinical symptoms, cognitive performance, and inflammatory markers were examined using both frequentist and Bayesian approaches.

**Results:** Five recurrent dynamic brain states were identified. Compared with controls, PCS participants showed reduced probability of occurrence and shorter lifetime of a visual/dorsal attention state, alongside increased probability of a limbic/default mode network (DMN) state. PCS was also characterised by reduced transitions between visual/dorsal attention and frontoparietal–DMN states, and increased transitions from somatomotor/visual states toward the limbic-DMN configuration. Exploratory analyses revealed that greater expression of the limbic-DMN state was negatively associated with global cognitive performance (MoCA) and positively associated with serum IL-1β levels.

**Conclusions:** PCS is associated with a reorganisation of intrinsic brain dynamics, marked by a shift from externally oriented attentional states toward limbic-DMN configurations. This imbalance may reflect reduced cognitive flexibility and increased sensitivity to interoceptive or affective signals. The observed links between dynamic brain-state expression, cognitive impairment, and peripheral inflammation support a systems-level mechanism through which immune signalling may influence neural function in PCS. Dynamic functional connectivity offers a sensitive framework for capturing these alterations and may inform future mechanistic models of post-viral dysfunction and therapeutic targeting.

**Highlights:**

- Post-COVID-19 Syndrome (PCS) is linked to altered intrinsic brain dynamics at rest.
- Patients show reduced engagement of visual and attentional brain states.
- Increased recruitment of limbic-DMN states suggests a shift toward more internally, emotionally charged focussed brain activity.
- Brain-state dynamics are putatively associated with global cognition and IL-1β.
- Reduced brain-state flexibility may underlie cognitive and emotional symptoms in PCS.

## Introduction

Post-COVID-19 Syndrome (PCS), also referred to as Long COVID, affects a considerable number of individuals after infection with SARS-CoV-2^1, 2^. Even months after the acute phase has resolved, many people continue to experience symptoms such as fatigue, poor concentration, slowed thinking, mood changes, and issues with memory^1–3^. These difficulties occur even in individuals who were not hospitalised and may persist despite normal findings on conventional brain imaging. The underlying causes of PCS are still being investigated, but current evidence suggests that the brain may be particularly vulnerable to subtle changes in how it functions and our different brain regions communicate with each other, especially in the context of inflammation^2, 4, 5^.

Resting-state functional MRI (rs-fMRI) has become a widely used tool for investigating large-scale brain network organisation in both health and disease^6^. By measuring spontaneous fluctuations in the Blood Oxygenation Level Dependent (BOLD) signal while participants are not engaged in any specific task, rs-fMRI reveals patterns of synchronised activity across different brain regions, thus uncovering the brain’s intrinsic functional architecture. These networks - such as the default mode, frontoparietal, limbic, and attention systems - are involved in diverse domains including emotion regulation, memory, self-reflection, and goal-directed cognition^6^. Notably, rs-fMRI measures have been shown to predict individual patterns of brain engagement during cognitive and affective tasks, offering insight into latent network dynamics that shape behaviour even at rest^6, 7^. Several studies using rs-fMRI have shown that PCS is associated with changes in connectivity within and between key large-scale brain networks^8–17^. In particular, altered connectivity has been observed in the default mode network (DMN), the salience network, and frontoparietal systems, which are responsible for executive control and attention^18, 19^ ^20^. These abnormalities often correlate with symptoms like fatigue and mental fog^14^. However, most of these studies have used “static” analyses that summarise brain functional connectivity as if it remained the same across the entire scan^21^. In reality, brain connectivity is constantly evolving^22^. Even at rest, the brain shifts from one functional state to another over the course of seconds. These states, which represent groups of brain regions whose activity is highly synchronised at a given moment in time, reflect distinct modes of neural processing^23^. The way these states emerge and take turns activating across time offers insights into the brain’s moment-to-moment responsiveness and its ability to flexibly adapt to changing demands ^22, 24, 25^.

In healthy individuals, the brain naturally moves between states related to attention, sensory awareness, emotion, and self-monitoring^23^. These states are temporary, and the brain usually does not get “stuck” in any one of them^22^. In psychiatric and neurological conditions such as depression, schizophrenia or body dysmorphic disorder, this balance is often disrupted. For example, some individuals spend more time in states related to internal thinking or negative emotion and explore fewer states related to attention and engagement with the external world^26–30^. Because individuals with PCS share many of these symptoms, it is reasonable to suspect that something similar may be happening in their brains. One method that allows us to investigate this in detail is called Leading Eigenvector Dynamics Analysis (LEiDA)^23, 31^. This technique examines moment-to-moment changes in how different brain regions synchronise their activity. LEiDA identifies a set of distinct brain states and shows how frequently each state occurs, how long it lasts, and how often the brain switches between them. Although LEiDA has not yet been widely used in PCS research, early connectivity studies suggest that individuals with PCS may show reduced network efficiency over time^14^, which in turn provides a potential backbone for less flexible brain dynamics.

In this context, we set out to use LEiDA to explore how large-scale brain states unfold over time in individuals with PCS during resting-state fMRI. Building on prior research linking PCS to cognitive and affective dysfunction^32, 33^ - as well as emerging evidence of altered network connectivity in fatigue-related and inflammatory conditions ^34^ - we formulated three core hypotheses. First, we hypothesised that individuals with PCS would spend more time in brain states dominated by the DMN and limbic regions. These areas are associated with internally directed cognition, self-referential thought, and affective processing, which may become predominant in the context of fatigue, emotional dysregulation, and withdrawal from external demands^19^. Second, we expected reduced occupancy of brain states engaging frontoparietal and dorsal attention networks, which are critical for sustained attention, goal-directed behaviour, and environmental monitoring^18, 35^. A diminished ability to access or maintain these externally focused states could help explain the common experiences of mental fog, distractibility, and slowed cognitive processing reported by PCS patients. Third, we hypothesised that individual differences in brain-state dynamics - specifically, reduced engagement of attention-related states and increased expression of DMN-limbic configurations - would be associated with clinical symptom severity, cognitive performance deficits (particularly in attention), and elevated peripheral markers of inflammation. Together, these hypotheses aimed to link brain dynamics with the neurocognitive and immunological signatures of PCS.

## Methods

### Study Design and Participants

As described elsewhere, this study employed a single-site observational case-control design. All participants, both PCS cases and recovered controls, were required to have had biologically confirmed SARS-CoV-2 infection at least three months prior to enrolment, consistent with the WHO clinical case definition of Post-COVID-19 condition^36^. Individuals in the PCS group were characterised by persistent symptoms lasting ≥3 months after infection, whereas recovered controls were required to be fully asymptomatic, with complete resolution of all post-COVID symptoms. We included individuals with mild acute COVID-19 only, to minimise potential confounding effects of hospitalisation, hypoxia, or severe systemic inflammation on brain perfusion and metabolism, and to ensure that any observed group differences were not attributable to the neurological sequelae of moderate–severe disease. All participants were required to have had biologically confirmed SARS-CoV-2 infection prior to enrolment. Confirmation was based on either a positive reverse-transcription polymerase chain reaction (RT-PCR) test from a nasopharyngeal swab or documented serological evidence of prior infection, in accordance with UK national testing guidelines at the time of acute illness. Participants with only clinically suspected but biologically unconfirmed COVID-19 were not included. Recovered healthy control participants were required to have a history of biologically confirmed SARS-CoV-2 infection, followed by complete resolution of all acute and post-acute symptoms. To minimize the risk of including individuals with subtle or subclinical post-COVID manifestations, control participants were screened using the same standardized clinical instruments applied to the PCS group, including the Fatigue Assessment Inventory. Only individuals scoring below established clinical thresholds, and reporting no persistent fatigue, cognitive complaints, affective symptoms, autonomic dysfunction, or post-exertional malaise at the time of assessment, were included as healthy controls. In addition, control participants underwent the same cognitive assessment battery and clinical interview as PCS participants, and individuals demonstrating clinically relevant deviations from normative performance or reporting residual symptoms were excluded. This stringent screening strategy ensured that the control group represented a recovered post-COVID population without ongoing symptomatology, rather than asymptomatic or mildly affected PCS cases.

We recruited participants through King’s College Hospital NHS Trust and surrounding community networks. Eligible individuals were aged 16–65 years, fluent in English, and capable of providing informed consent. We excluded individuals with a history of major neurological or psychiatric disorders, current substance misuse, systemic or central nervous system inflammatory conditions, chronic respiratory or cardiac disease, recent vaccination (within two weeks), pregnancy or breastfeeding, BMI >30, or current use of immunomodulatory or psychoactive medications. We used the global severity score of Fatigue Assessment Inventory (FAI) to pre-screen participants for symptom severity ^37^. Individuals scoring greater than 4 on the global fatigue severity scale were classified into the PCS group with persistent fatigue, while those scoring 4 or below were included as recovered controls. The threshold of > 4 was selected to ensure inclusion of individuals with clinically meaningful levels of fatigue, consistent with prior applications of the FAI in post-viral and chronic fatigue contexts^37^. Groups were balanced for age, sex, BMI, and acute COVID-19 severity. The study received ethical approval from the UK Health Research Authority (IRAS ID: 308661) and was conducted in accordance with the Declaration of Helsinki and Good Clinical Practice (GCP) guidelines. All participants provided written informed consent prior to study procedures.

We conducted an a priori power analysis using G*Power (version 3.1) to estimate the required sample size for detecting group differences in dynamic functional connectivity metrics using analysis of covariance (ANCOVA). Assuming a medium to large effect size (Cohen’s f=0.30, equivalent to ηp^2^≈0.08), alpha = 0.05, power (1 − β) = 0.80, and three covariates (age, sex, and handedness), the estimated sample size required per group was 19. Our sample of 20 participants per group therefore provides adequate power to detect effects in this range.

### Procedures

Each participant completed a single study visit that included clinical assessments, venous blood collection, and a multimodal MRI scan. Clinical assessments included medical and psychiatric history, physical examination, vital signs, and a structured battery of standardized questionnaires. Fatigue was assessed using both the full Fatigue Assessment Inventory^37^ and the Multidimensional Fatigue Inventory^38^ to differentiate between mental and physical fatigue. Symptoms of autonomic dysfunction were evaluated using the Composite Autonomic Symptom Score (COMPASS-31)^39^, while mood and anxiety were measured using the Hospital Anxiety and Depression Scale (HADS)^40^. Sleep quality was assessed with the Pittsburgh Sleep Quality Index^41^, and respiratory symptoms were rated using the MRC Dyspnoea Scale^42^. Additional measures included the DePaul Symptom Questionnaire^43^, the American College of Rheumatology (ACR) fibromyalgia criteria^44^, the PAIN Detect questionnaire^45^, and Montreal Cognitive Assessment^46^. Most self-report questionnaires were completed remotely via the Qualtrics platform within 72 hours of the in-person visit. All participants underwent a structured diagnostic assessment using the Mini International Neuropsychiatric Interview (MINI) to exclude current or past major psychiatric disorders, including mood, anxiety, psychotic, and substance use disorders.

### Blood analysis

Venous blood samples (up to 30 mL) were collected on the day of imaging. Standard hematological and biochemical panels - including erythrocyte sedimentation rate (ESR), C-reactive protein (CRP), thyroid-stimulating hormone (TSH), free thyroxine (T4), and liver function tests - were processed by Synnovis (Viapath) at King’s College Hospital NHS Foundation Trust. Serum samples were frozen at −80°C and later assayed for cytokines and glial markers using electrochemiluminescence immunoassays (Meso Scale Discovery, Rockville, MD, USA). The cytokine panel included interleukin-1 beta (IL-1β), interleukin-2 (IL-2), interleukin-4 (IL-4), interleukin-6 (IL-6), interleukin-8 (IL-8), interleukin-10 (IL-10), interleukin-12p70 (IL-12p70), interleukin-13 (IL-13), tumor necrosis factor-alpha (TNF-α), and interferon-gamma (IFN-γ). Glial markers included glial fibrillary acidic protein (GFAP), measured with the ultrasensitive kit, and S100β. Assays were performed in duplicate. Values below the detection limit were discarded. This led us to discard all quantifications of IL-2, IL-4 and IL-12p70. Outliers greater than three standard deviations from the group mean were winsorized. No analyte had more than 10% missing data, therefore all data were included.

### Cognitron

Cognitive performance was assessed online using a battery of computerized tasks from the Cognitron platform, covering delayed verbal memory, working memory, executive function, attention, motor coordination, and spatial manipulation^33^. Tasks included Verbal Analogies (abstract reasoning and semantic integration), Immediate and Delayed Prospective Memory (episodic memory and planning), 2D Spatial Manipulation (visuospatial working memory and mental rotation), Motor Control (sensorimotor coordination and reaction consistency), Spotter (vigilance and sustained attention), and Block Reasoning (fluid intelligence and rule-based problem solving). Each task involves trial-level response collection, capturing both accuracy and reaction time. Performance metrics were transformed into standardized deviation-from-expected (DFE) scores using normative models derived from a large independent sample (n > 20,000), adjusted for age, sex, handedness, and harmonized ethnicity. DFE scores represent standardized residuals (observed minus expected performance divided by normative standard deviation) and were computed separately for accuracy and reaction time.

## Neuroimaging Data Acquisition and Processing

### MRI Acquisition

We acquired MRI data using a 3T GE Healthcare MR750 scanner (GE Medical Systems, Milwaukee, WI), equipped with a 32-channel receive-only head coil. The imaging protocol was part of the larger FALCS neuroimaging study and included high-resolution anatomical scans, quantitative BOLD sequences, and cerebral perfusion imaging. For the purposes of this study, we focused exclusively on resting-state functional MRI (rs-fMRI) data to assess dynamic brain connectivity.

T1-weighted structural images were acquired using a 3D magnetisation-prepared rapid acquisition gradient echo (MPRAGE) sequence with the following parameters: repetition time (TR) = 2300 ms, echo time (TE) = 2.98 ms, inversion time (TI) = 900 ms, and voxel size = 1 mm³ isotropic. These anatomical images served as the reference for spatial registration and normalisation.

Resting-state fMRI data were acquired using a multi-echo echo-planar imaging (ME-EPI) sequence with full k-space sampling at each echo. Three echoes were collected per volume with echo times (TEs) of 13 ms, 28 ms, and 44 ms. The repetition time (TR) was 2000 ms, and a total of 240 volumes were acquired over 8 minutes, following four initial dummy scans to allow for signal stabilization. Imaging covered the whole brain with 32 near-axial slices oriented along the anterior commissure–posterior commissure (AC–PC) line, each 3 mm thick with a 1 mm inter-slice gap. The field of view (FOV) was 240 mm, and the flip angle was set to 80 degrees. During scanning, participants were instructed to remain still, keep their eyes open, and fixate on a central crosshair.

### Preprocessing

We conducted preprocessing using FSL (version 6.0) and tedana.py (version 0.0.10; https://tedana.readthedocs.io/en/latest). The first step involved registering the anatomical MPRAGE image to Montreal Neurological Institute (MNI) space. Functional volumes from the first echo of each rs-fMRI time series were aligned to the first volume using mcflirt in FSL. The resulting transformation matrices were then applied to the remaining echoes, ensuring consistent alignment across all echoes. We then performed T2*-weighted optimal combination of the echoes to enhance BOLD contrast, and denoised the data using multi-echo independent component analysis (ME-ICA) implemented in tedana.py^47, 48^. This method decomposes the data into independent components and classifies them as BOLD or non-BOLD based on their TE-dependence. Components identified as non-BOLD - typically reflecting motion, physiological noise, or scanner-related artefacts - were regressed out of the optimally combined data, yielding a denoised multi-echo BOLD dataset. The denoised time series were then registered to the corresponding MPRAGE structural image using epi_reg and subsequently normalised to MNI space using the previously computed anatomical transformations. All functional images were resampled to 2 mm³ isotropic resolution. To enhance spatial comparability and reduce high-frequency noise, the data were smoothed using a Gaussian kernel with 6 mm full-width at half-maximum (FWHM). We applied high-pass temporal filtering with a cut-off frequency of 0.01 Hz to remove low-frequency drifts and scanner instabilities. Finally, to account for residual non-neuronal sources of variance, we regressed out the average signal from white matter (WM) and cerebrospinal fluid (CSF). WM and CSF masks were defined in MNI space using the MNI152 standard template and applied to each participant’s preprocessed data. The extracted mean WM and CSF signals were regressed out using fsl_glm to reduce physiological and scanner-related noise, such as cardiac pulsatility and respiratory fluctuations. This preprocessing pipeline yielded denoised, anatomically aligned, and temporally filtered functional data optimised for subsequent dynamic connectivity analyses^49^.

### Quality Control

All structural and functional images were visually inspected for artefacts, motion, and co-registration accuracy at multiple stages of preprocessing. We verified alignment between functional and anatomical images following each registration step, including echo combination, denoising, and MNI normalisation. For the rs-fMRI data, we monitored framewise displacement and excluded any datasets exceeding a mean framewise displacement of 0.5 mm or containing more than 10% of volumes with motion spikes above 0.75 mm. No participants exceeded these thresholds. Independent component classification by ME-ICA was visually examined in a subset of participants to confirm the effective separation of BOLD and non-BOLD components. Additionally, all white matter and CSF masks used for nuisance regression were manually checked for correct anatomical placement in MNI space. These quality control procedures ensured robust and reliable inputs for the dynamic functional connectivity analyses.

### LEiDA Analyses

We performed Leading Eigenvector Dynamics Analysis (LEiDA) in MATLAB R2020a (MathWorks, Natick, MA, USA) using scripts adapted from *Cabral et al. (2017)* [https://github.com/juanitacabral/LEiDA]. This method tracks how synchrony between different brain regions evolves over time by analysing the timing of their BOLD signal fluctuations.

We first extracted BOLD time series from 105 cortical and subcortical regions of interest (ROIs) using the Harvard–Oxford atlas available from the CONN toolbox^50^. Cerebellar ROIs were excluded due to their absence in the Yeo 7-network parcellation. Each ROI time series was then Hilbert-transformed to derive the instantaneous phase in the complex plane (Figure 1A). At each timepoint, we computed a dynamic phase-locking matrix dPL(t), quantifying pairwise phase synchrony between regions (Figure 1B). We extracted the leading eigenvector V□(t) from each dPL(t) to capture the dominant pattern of whole-brain synchrony at each timepoint. We then concatenated V□(t) across all participants and applied k-means clustering to identify recurrent brain states (Figure 1C). We tested clustering solutions from k = 5 to k = 10 and selected the optimal number of states (k = 5) using the Dunn index, which maximises between-cluster separation and within-cluster homogeneity.

**Figure 1.**
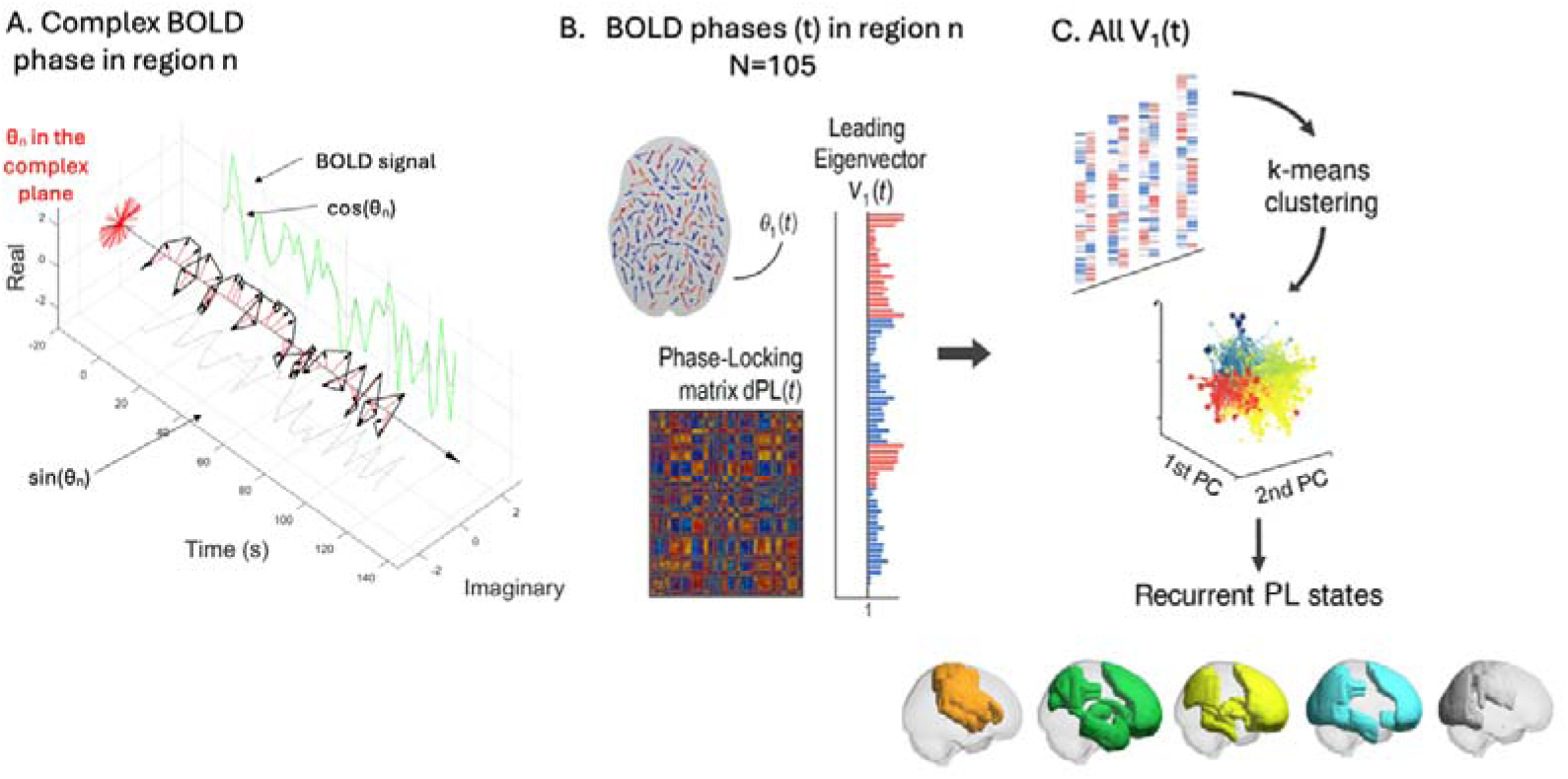
Schematic overview of the LEiDA pipeline for identifying recurrent brain states. (A) For each region n, the BOLD signal is bandpass-filtered and transformed into its complex representation to extract the instantaneous phase θn(t) via the Hilbert transform. (B) At each timepoint t, the phases across 105 regions are used to compute a dynamic phase-locking matrix dPL(t), from which the leading eigenvector V1(t) is extracted. V1(t) captures the dominant pattern of phase synchrony across the brain at that time. (C) All leading eigenvectors V1(t) across participants and timepoints are concatenated and submitted to k-means clustering, identifying a set of recurrent phase-locking (PL) states. Each state reflects a spatially distinct and temporally recurring pattern of functional connectivity.

For each identified state, we computed three dynamic metrics per participant: (1) probability of occurrence (proportion of time spent in the state), (2) average lifetime (number of consecutive timepoints within the state), and (3) pairwise transition probabilities between states.

To interpret the functional relevance of each state, we calculated Spearman correlations between the spatial topography of each state’s centroid (Vk) and the canonical Yeo 7-network maps^51^. These Yeo networks include the visual, somatomotor, dorsal attention, ventral attention, limbic, frontoparietal, and default mode networks. Significance was defined using a Bonferroni-corrected threshold of p < 0.05/7 (0.007).

### Statistical Analysis

We compared group differences in LEiDA-derived dynamic metrics using analysis of covariance (ANCOVA), controlling for age, sex, and handedness. For each state, we examined between-group differences in probability of occurrence, lifetime, and transition probabilities. For completeness, we supplemented frequentist statistics with their respective Bayesian counterparts to quantify evidence strength. Bayes factors (BF□□) above 1, 3 and 10 were interpreted as anecdotal, moderate, and strong evidence for a given effect, respectively.

We also performed exploratory correlation analyses between dynamic metrics and behavioural or biological measures, including symptom severity (e.g., fatigue, anxiety), cognitive performance (e.g., sustained attention), and circulating inflammatory markers (e.g., interleukin-6). We used partial Spearman rank correlations, which accounted for age, gender, handedness and group.

## Results

### Sociodemographics and clinical characterisation

Results have been first reported elsewhere. Here we describe the main findings that might be relevant to interpret the new neuroimaging findings derived from the use of functional BOLD resting-state imaging. The groups were matched for age, sex, ethnicity, and body mass index (BMI), with no significant differences observed across these variables (all p > 0.05; Table 1). However, groups differed significantly in years of education, with HC participants reporting higher educational attainment. PCS participants also had a significantly longer duration since first SARS-CoV-2 infection, and were significantly less likely to have been vaccinated prior to infection. Clinically, individuals with PCS reported significantly greater symptom burden across multiple domains. These included fatigue (both visual analogue scale and multidimensional components), sleep disturbance (PSQI), post-exertional malaise, autonomic dysfunction (COMPASS-31), affective symptoms (HADS depression/anxiety), PTSD-related symptoms, and musculoskeletal complaints (WPI and SS scores). Notably, 40% of PCS participants met criteria for fibromyalgia, compared to none in the HC group. These differences were supported by both frequentist (p < 0.05) and Bayesian inference, with many outcomes yielding strong to extreme evidence in favour of group separation (BF□□ > 10), particularly for fatigue, post-exertional malaise, and pain sensitivity (Supplementary Table S1).

**Table 1.** Sociodemographic and clinical characteristics of PCS and healthy control participants. Group comparisons of demographic and COVID-19-related variables between individuals with Post-COVID-19 Syndrome (PCS) and matched healthy controls (HC). Data are shown as mean (standard deviation) or counts. Between-group differences were assessed using independent samples t-tests or chi-squared tests as appropriate. Bayes factors (BF□□) quantify evidence for group differences; values >1 indicate increasing support for the alternative hypothesis. Asterisks (*) denote statistically significant results at p < 0.05 or BF□□ > 1.

| Variable | PCS Mean (std) | HC Mean (std) | Statistics | p | $B_{10}$ |
| --- | --- | --- | --- | --- | --- |
| Age | 40.6 (10.9) | 40.0 (10.4) | $T(38)=0.208$ | 0.837 | 0.312 |
| Gender | 18F/2M | 17F/3M | $\chi^2(1)=0.229$ | 0.663 | N/A |
| Ethnicity | White: 18<br>Mixed: 0<br>Asian: 1<br>Black: 0<br>South Asian: 1<br>Hispanic: 0 | White: 15<br>Mixed: 1<br>Asian: 1<br>Black: 2<br>South Asian: 0<br>Hispanic: 1 | $\chi^2(5)=5.273$ | 0.384 | N/A |
| BMI | 23.8 (3.44) | 24.4 (3.84) | $T(38)=-0.55$ | 0.582 | 0.346 |
| SPO<sub>2</sub> (%) | 99.105 (1.524) | 98.842 (1.344) | $T(36)=0.565$ | 0.576 | 0.357 |
| Years of education | 17.30 (2.94) | 19.35 (3.13) | $T(38)=-2.13$ | 0.039* | 1.845 |
| MRC Dyspnea scale | 0: 45%<br>1: 45%<br>2: 10%<br>3: 0%<br>4: 0% | 0: 95%<br>1: 5%<br>2: 0%<br>3: 0%<br>4: 0% | $\chi^2(2)=11.971$ | 0.003* | N/A |
| Time since first infection (months) | 23.0 (9.17) | 15.7 (11.5) | $T(38)=2.21$ | 0.033* | 0.749 |
| **Vaccination prior infection** | Yes: 4<br>No: 16 | Yes: 14<br>No: 6 | $\chi^2(1)=10.101$ | 0.001* | N/A |

### Clinical routine blood tests

Group comparisons of routine blood parameters revealed no major differences between PCS and HC participants (Supplementary Table S2). However, PCS participants showed numerically higher creatinine and Gamma-GT levels, with corresponding Bayes factors (BF□□ = 1.232 and 1.735, respectively) providing anecdotal to moderate evidence for group differences. Platelet count also trended higher in the PCS group (p = 0.08; BF□□ = 1.123). Other metabolic, inflammatory, and hematological markers - including C-reactive protein, white and red cell counts, hemoglobin, and thyroid function - did not differ significantly between groups.

### Serum cytokines and glial markers

No statistically significant differences were observed in circulating levels of IFN-γ, IL-1β, IL-6, IL-8, IL-10, IL-13, TNF-α or GFAP (Supplementary Table S3). However, TNF-α (p = 0.116; BF□□ = 1.094) and S100β (p = 0.149; BF□□ = 1.025) showed suggestive evidence of elevation in the PCS group, consistent with mild glial or immune activation.

### Cognitive performance

Cognitive testing using the Cognitron battery revealed subtle but functionally relevant differences between groups (Supplementary Table S4). PCS participants showed significantly poorer performance on the delayed object memory task (RT_DFE; p = 0.02; BF□□ = 2.653), as well as trend-level impairments in the Lead Balloon task (abstract reasoning; p = 0.06; BF□□ = 1.330) and vigilance (Spotter task RT; p = 0.06; BF□□ = 1.303). Other domains - including motor control, 2D spatial manipulation, and verbal analogies - did not show significant differences. Overall, Bayesian analysis provided moderate evidence for reduced memory and attentional function in PCS.

### Identification and characterisation of optimal number of dynamic brain states

Using the Dunn index, we identified five recurrent brain states maximising within-state homogeneity and between-state separability. Each state exhibited a distinct spatial configuration of phase synchrony, mapped onto canonical Yeo networks. State 1 primarily involved visual and dorsal attention networks; State 2 showed strong expression in somatomotor and ventral attention regions; State 3 involved mainly visual networks; State 4 was characterised by coactivation of frontoparietal and default mode network (DMN) regions; and State 5 was dominated by limbic and DMN components (Figure 2 and Table 2).

**Figure 2.**
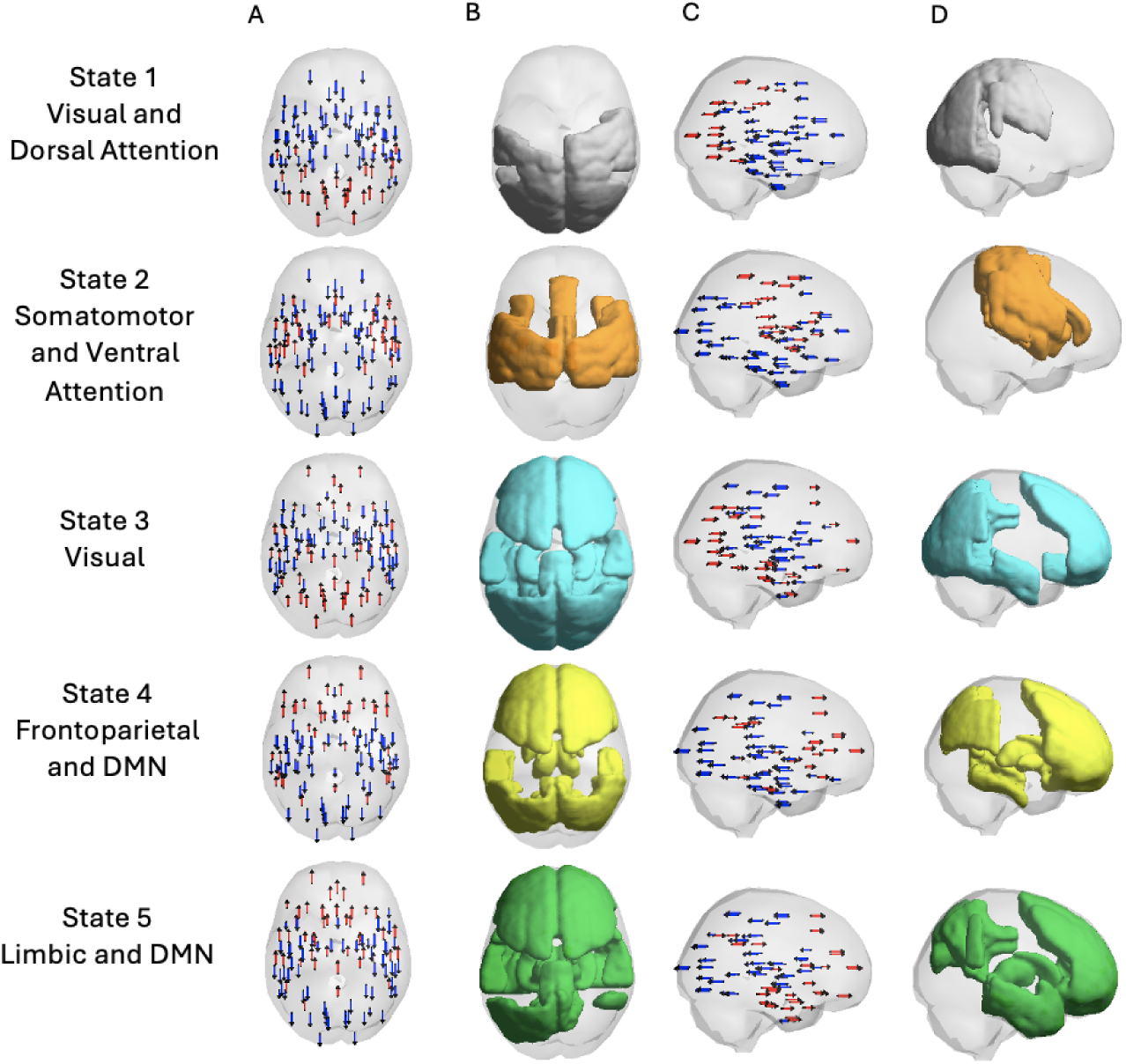
Recurrent Large-Scale Brain States Identified with LEiDA. This figure depicts the five dynamic functional brain states extracted using Leading Eigenvector Dynamics Analysis (LEiDA) from resting-state fMRI data. Each row represents one recurrent state, characterized by its dominant spatial organization and network affiliations: (1) Visual and Dorsal Attention, (2) Somatomotor and Ventral Attention, (3) Visual, (4) Frontoparietal and Default Mode Network (DMN), and (5) Limbic and DMN. Column A shows the leading eigenvector phase alignment patterns in axial view, with blue and red arrows indicating opposite phase relationships across regions. Column B highlights the brain areas with the highest contributions to each state, mapped onto inflated cortical surfaces. Column C displays lateral views of the eigenvector vectors to illustrate the directionality and distribution of phase coupling across the cortex. Column D presents an alternative cortical view of the same spatial distribution shown in Column B to aid anatomical interpretation.

**Table 2.** Functional network composition of dynamic brain states. Each row shows the Spearman correlation (ρ) between the spatial configuration of a given dynamic brain state (States 1–5) and the canonical Yeo 7-network parcellation: Visual, Somatomotor Network (SMN), Dorsal Attention (DA), Ventral Attention (VA), Limbic, Frontoparietal Network (FPN), and Default Mode Network (DMN). Significant correlations after bonferroni correction are marked with an asterisk (*). Values in parentheses indicate uncorrected p-values. State labels reflect the dominant functional networks positively associated with each state.

| State number | Labels | Visual | SMN | DA | VA | Limbic | FPN | DMN |
| --- | --- | --- | --- | --- | --- | --- | --- | --- |
| 1 | Visual/DA | 0.600*<br>($<0.001$) | 0.067<br>(0.494) | 0.484*<br>($<0.001$) | 0.062<br>(0.530) | -0.359*<br>($<0.001$) | 0.078<br>(0.429) | -0.338*<br>($<0.001$) |
| 2 | SMN/VA | -0.480*<br>($<0.001$) | 0.610*<br>($<0.001$) | 0.110<br>(0.264) | 0.529*<br>($<0.001$) | -0.304*<br>(0.002) | 0.085<br>(0.388) | -0.328*<br>($<0.001$) |
| 3 | Visual | 0.704*<br>($<0.001$) | -0.598*<br>($<0.001$) | 0.124<br>(0.208) | -0.540*<br>($<0.001$) | 0.134<br>(0.171) | -0.033<br>(0.735) | 0.162<br>(0.098) |
| 4 | FPN/DMN | -0.299*<br>(0.002) | -0.382*<br>($<0.001$) | 0.315*<br>(0.001) | 0.211<br>(0.030) | -0.013<br>(0.897) | 0.733*<br>($<0.001$) | 0.442*<br>($<0.001$) |
| 5 | Limbic/DMN | -0.146<br>(0.137) | -0.465*<br>($<0.001$) | -0.042<br>(0.672) | -0.294*<br>(0.002) | 0.411*<br>($<0.001$) | 0.242<br>(0.013) | 0.594*<br>($<0.001$) |

### Group Differences in Brain-State Dynamics

We conducted ANCOVAs controlling for age, sex, and handedness to compare dynamic metrics between individuals with PCS and healthy controls. Participants with PCS exhibited a significantly lower probability of occurrence and shorter lifetime in State 1. Conversely, they showed a higher probability of occurrence for State 5. No significant group differences were found in States 2–4 after correcting for multiple comparisons. Bayesian counterparts generally supported these observations (Table 3 and Figure 3).

**Figure 3.**
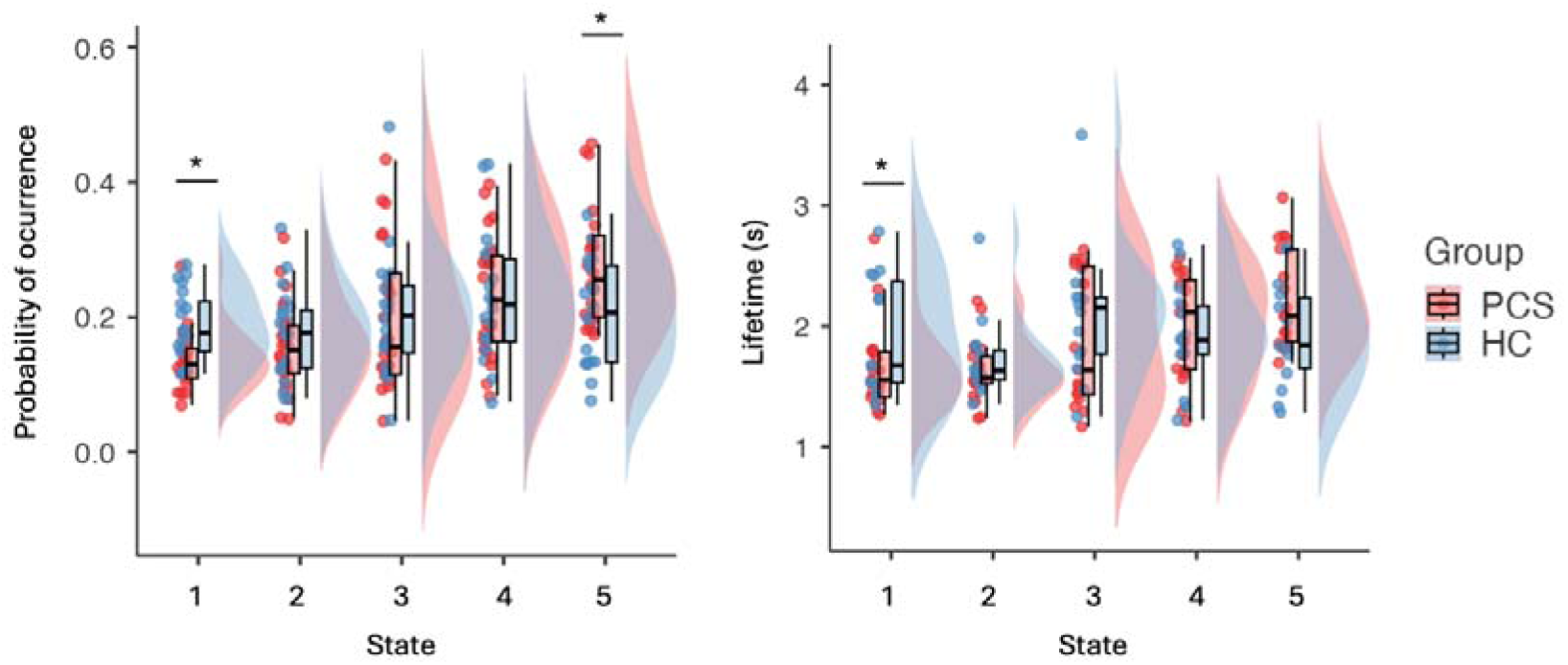
Group differences in dynamic brain-state metrics between PCS and healthy controls. Violin plots show (left) the probability of occurrence and (right) the average lifetime (in seconds) for each of the five recurrent phase-locking states identified using LEiDA. Individuals with Post-COVID-19 Syndrome (PCS; red) exhibited a significantly lower probability of occurrence and shorter lifetime in State 1 (visual/dorsal attention) and a significantly higher probability of occurrence in State 5 (limbic/DMN), compared to healthy controls (HC; blue). *p < .05, ANCOVA controlling for age, sex, and handedness; error bars represent interquartile range with median.

**Table 3.** Group comparisons of dynamic brain-state metrics between PCS and healthy controls. Mean (standard deviation) values are shown for the probability of occurrence and average lifetime (in seconds) for each of the five LEiDA-derived brain states. Statistical results reflect ANCOVAs controlling for age, sex, and handedness. *p < .05. BF□□ refers to the Bayes Factor in favour of the alternative hypothesis (PCS ≠ HC), with values >3 indicating moderate and >10 strong evidence. Significant differences were observed for States 1 and 5 in probability of occurrence, and for State 1 in lifetime.

| Metric | State | PCS Mean (SD) | HC Mean (SD) | Statistics | P-value | $B_{10}$ |
| --- | --- | --- | --- | --- | --- | --- |
| Probability | 1 | 0.136<br>(0.045) | 0.185<br>(0.053) | $F(1,35)=9.443$ | 0.004* | 13.59* |
|  | 2 | 0.157<br>(0.069) | 0.175<br>(0.066) | F(1,35)=0.796 | 0.378 | 0.443 |
|  | 3 | 0.198<br>(0.111) | 0.205<br>(0.093) | F(1,35)=0.045 | 0.842 | 0.316 |
|  | 4 | 0.233<br>(0.095) | 0.228<br>(0.0941) | F(1,35)=0.084 | 0.774 | 0.319 |
|  | 5 | 0.277<br>(0.092) | 0.207<br>(0.081) | F(1,35)=5.862 | 0.021* | 3.57* |
| **Lifetime** | 1 | 1.66<br>(0.363) | 1.96<br>(0.466) | F(1,35)=5.031 | 0.032* | 2.39* |
|  | 2 | 1.67<br>(0.306) | 1.73<br>(0.318) | F(1,35)=0.208 | 0.652 | 0.359 |
|  | 3 | 1.86<br>(0.553) | 2.06<br>(0.534) | F(1,35)=1.347 | 0.254 | 0.545 |
|  | 4 | 2.04<br>(0.510) | 1.96<br>(0.398) | F(1,35)=0.389 | 0.537 | 0.358 |
|  | 5 | 2.23<br>(0.409) | 2.08<br>(0.620) | F(1,35)=0.631 | 0.433 | 0.417 |

### Transition Probabilities

We analysed state-to-state transition probabilities to assess how participants moved between dynamic configurations. PCS participants showed reduced transitions from State 4 to State 1 and from State 1 to State 4. Conversely, transitions from States 2 and 3 to State 5 were significantly increased in the PCS group, indicating a tendency to dwell in internally focused configurations (Figure 4 and Table 4).

**Figure 4.**
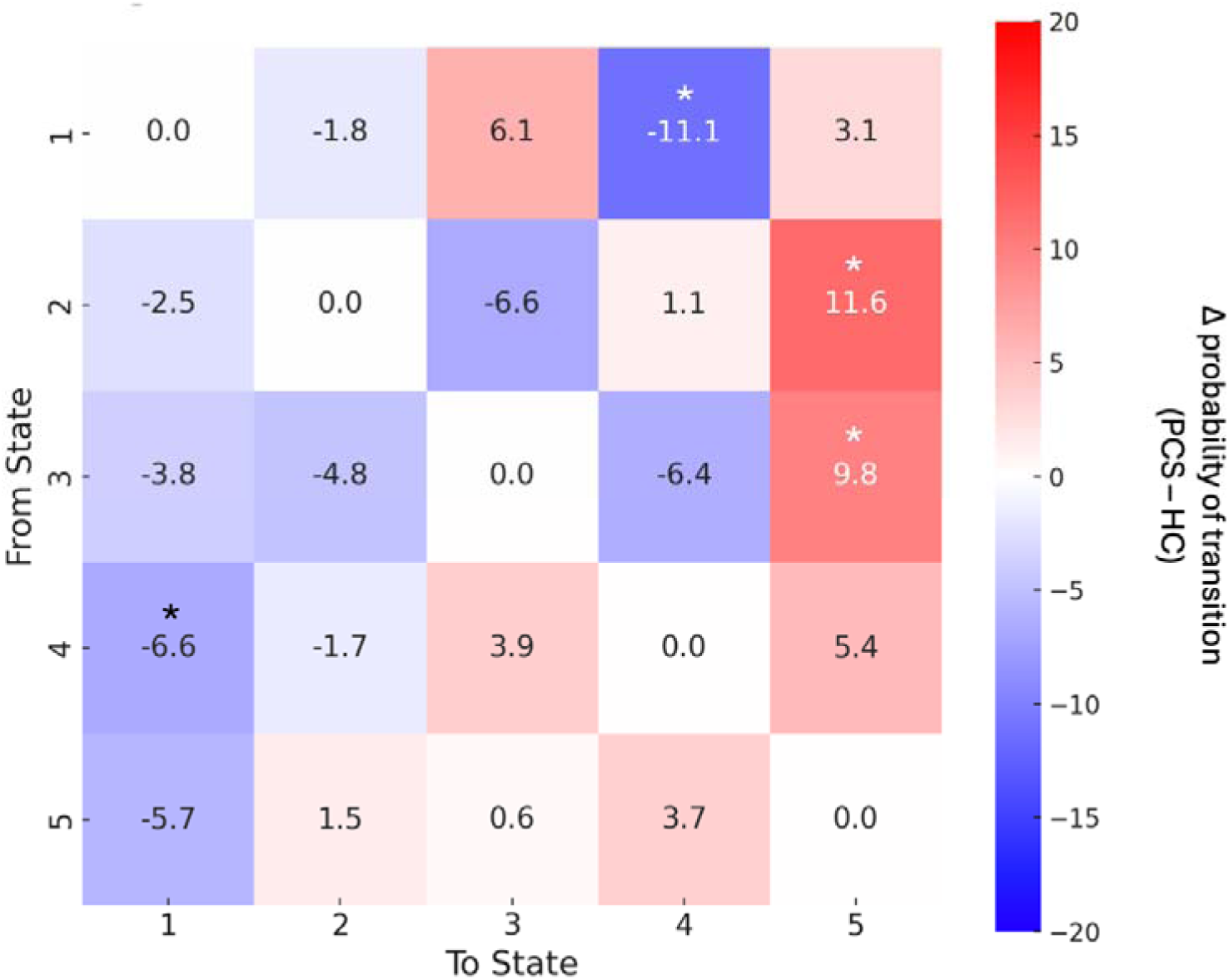
Group differences in transition probabilities between dynamic brain states. Heatmap depicts the difference in transition probabilities between PCS and healthy control groups (Δ = PCS − HC) for each pair of phase-locking states. Positive values (red) indicate increased transition probability in PCS; negative values (blue) indicate decreased transition probability. PCS participants showed significantly reduced transitions between State 4 (frontoparietal/DMN) and State 1 (visual/dorsal attention), and significantly increased transitions from State 2 (somatomotor/ventral attention) and State 3 (visual) into State 5 (limbic/DMN). Asterisks denote group differences significant at *p < .05, ANCOVA controlling for age, sex, and handedness.

**Table 4.** Group differences in transition probabilities between dynamic brain states. Mean transition probabilities (± standard deviation) are shown for all possible pairwise transitions between the five dynamic brain states, separately for the PCS and healthy control (HC) groups. Group comparisons were conducted using ANCOVAs controlling for age, sex, and handedness, with corresponding p-values and Bayes Factors (BFlJlJ) reported.

| | Group | 1 → 2 | 1 → 3 | 1 → 4 | 1 → 5 | 2 → 1 | 2 → 3 | 2 → 4 | 2 → 5 | 3 → 1 | 3 → 2 | 3 → 4 | 3 → 5 | 4 → 1 | 4 → 2 | 4 → 3 | 4 → 5 | 5 → 1 | 5 → 2 | 5 → 3 | 5 → 4 |
| --- | --- | --- | --- | --- | --- | --- | --- | --- | --- | --- | --- | --- | --- | --- | --- | --- | --- | --- | --- | --- | --- |
| Mean | PCS | 0.193 | 0.301 | 0.268 | 0.277 | 0.207 | 0.253 | 0.257 | 0.355 | 0.205 | 0.218 | 0.235 | 0.342 | 0.177 | 0.225 | 0.252 | 0.356 | 0.190 | 0.248 | 0.239 | 0.345 |
|  | HC | 0.211 | 0.240 | 0.379 | 0.246 | 0.232 | 0.319 | 0.246 | 0.239 | 0.243 | 0.266 | 0.259 | 0.244 | 0.243 | 0.242 | 0.213 | 0.302 | 0.247 | 0.233 | 0.223 | 0.308 |
| SD | PCS | 0.105 | 0.146 | 0.120 | 0.128 | 0.145 | 0.139 | 0.168 | 0.151 | 0.0828 | 0.104 | 0.102 | 0.124 | 0.0728 | 0.0994 | 0.135 | 0.164 | 0.0786 | 0.104 | 0.111 | 0.124 |
|  | HC | 0.124 | 0.110 | 0.163 | 0.129 | 0.123 | 0.169 | 0.101 | 0.115 | 0.0915 | 0.147 | 0.102 | 0.134 | 0.102 | 0.121 | 0.120 | 0.105 | 0.147 | 0.105 | 0.0904 | 0.119 |
| Frequentist | F(1,32) | 0.452 | 1.411 | 7.219 | 0.553 | 0.269 | 1.820 | 0.122 | 6.287 | 1.747 | 1.499 | 0.498 | 5.321 | 4.969 | 0.297 | 1.100 | 1.359 | 2.206 | 0.131 | 0.362 | 0.878 |
|  | P-value | 0.506 | 0.244 | 0.011 | 0.462 | 0.589 | 0.186 | 0.729 | 0.018 | 0.195 | 0.229 | 0.485 | 0.027 | 0.032 | 0.589 | 0.301 | 0.252 | 0.147 | 0.719 | 0.551 | 0.355 |
| Bayesian | B<sub>10</sub> | 0.380 | 0.616 | 4.96 | 0.402 | 0.354 | 0.663 | 0.331 | 4.11 | 0.659 | 0.583 | 0.382 | 2.82 | 2.40 | 0.350 | 0.480 | 0.566 | 0.770 | 0.327 | 0.347 | 0.435 |

### Exploratory correlations between brain dynamics and clinical, blood and cognitive parameters

We conducted exploratory correlation analyses restricted to dynamic brain-state metrics that showed significant group differences - namely, the probability of occurrence and lifetime of State 1 (visual/dorsal attention) and State 5 (limbic/default mode network). The probability of occurrence of State 5 was negatively correlated with global cognitive performance on the MoCA (Spearman’s ρ = –0.43, p = 0.015) and with total serum protein levels (ρ = –0.37, p = 0.038), and positively correlated with circulating interleukin-1β (IL-1β; ρ = 0.39, p = 0.027), even though none of these correlations survived FDR correction for the number of parameters tested. In addition, the probability of occurrence of State 1 was positively associated with unrefreshing sleep severity (ρ = 0.34, p = 0.047). None of the remaining correlations reached significance (Supplementary Table S5).

## Discussion

This study provides novel evidence that PCS is associated with altered intrinsic brain dynamics at rest, particularly in large-scale networks supporting attention, sensory integration, and emotion regulation. Using Leading Eigenvector Dynamics Analysis (LEiDA), we identified five recurrent phase-locking states and showed that PCS participants spent significantly less time in an externally oriented attentional state (State 1: visual/dorsal attention) and more time in an internally affect-focused state (State 5: limbic/default mode network). These alterations were accompanied by reduced transitions between cognitively adaptive states and increased transitions into limbic-dominated configurations. Together, our findings suggest that PCS may involve a shift in baseline brain function away from goal-directed engagement and toward internally driven, potentially affectively charged modes of processing, an imbalance that may offer a fresh brain systems-level perspective on the mechanisms underlying key cognitive and emotional symptoms in PCS.

Our findings resonate with previous studies showing that healthy cognition depends not only on the strength of functional connectivity but also on the dynamic reconfiguration of brain networks over time^24^. In this context, brain-state flexibility - the ability to rapidly shift between large-scale functional configurations - has emerged as a key systems-level property supporting attention, working memory, and emotional control^24, 25^. Prior work using LEiDA and other dynamic functional connectivity methods has demonstrated that reduced variability or diminished access to specific dynamic states is associated with clinical conditions marked by fatigue, attentional deficits, and mood disturbances, such as major depression^26–28^. Our results suggest that PCS may share this broader signature of dynamic dysregulation, positioning it within a transdiagnostic framework of altered brain-state transitions.

The reduced probability and lifetime in State 1 among PCS participants is particularly compelling. This state overlapped with dorsal attention and visual networks systems essential for externally directed cognitive control^18, 52^. A reduced ability to enter or maintain this state suggests that patients may struggle to sustain attention or shift efficiently between perceptual inputs and task demands^53^. Importantly, the increased probability of occurrence of State 5, which involved the limbic system and DMN, indicates an imbalance rather than uniform suppression of dynamics. The limbic-DMN configuration is typically associated with self-referential thought, emotional salience, and interoceptive monitoring^35, 54^. Its overexpression may reflect a pathological bias toward internally focused processing, consistent with reports of emotional dysregulation, rumination, and heightened somatic awareness in PCS^19, 55, 56^.

Altered transition probabilities further reinforce the hypothesis of reduced dynamic repertoire and functional rigidity^57, 58^. In controls, more frequent reciprocal transitions between State 1 and the frontoparietal/DMN state (State 4) may reflect flexible toggling between goal-directed and introspective modes, akin to previously described metastable switching in healthy cognition^59, 60^. In contrast, PCS participants exhibited attenuated transitions between these states and increased transitions from somatomotor and visual states (States 2 and 3) into the limbic-DMN state (State 5). This asymmetric dynamic trajectory may reflect a form of neural inertia or “gravitational pull” into internally focused, affectively loaded brain modes, echoing prior findings in mood disorders and inflammatory syndromes^61, 62,20, 63^

While our findings reveal robust differences in the dynamic organisation of intrinsic brain activity between individuals with PCS and healthy controls, we emphasise the interpretive limitations of resting-state fMRI. The method captures large-scale patterns of synchronised neural activity but does not permit definitive conclusions about causality or mechanism. Interpreting these patterns functionally risks reverse inference, especially given that most cognitive or emotional processes map onto multiple, overlapping neural configurations rather than one-to-one brain-function relationships. By reframing our results through a transdiagnostic lens - where similar alterations in brain-state flexibility have been observed in conditions such as depression we align with the descriptive strengths of the LEiDA methodology while avoiding speculative or reductive claims. At the same time, we wish to avoid overcompensating in the opposite direction. Emphasising only peripheral or structural explanations risks overlooking more subtle cognitive and emotional contributions to symptom expression - particularly alterations in attention, emotional salience attribution, and interoceptive monitoring. These mechanisms are not mutually exclusive with dysautonomia or inflammation, but may in fact co-express and interact dynamically. Clinically, many individuals with persistent cognitive symptoms following COVID-19 infection present with features closely aligned with functional cognitive disorders - an area still under-recognised by many clinicians and neuroscientists. Contemporary theoretical frameworks, such as those drawing from predictive processing and ‘constructed emotion’ models, suggest that symptom persistence may reflect impaired updating of internal models about bodily and cognitive states. This can result in ‘symptom construction’ that continues beyond the resolution of initial peripheral or immune triggers, as seen in other conditions where inflammation-related delirium or fatigue transitions into persistent brain fog.

Although our study was not designed to explicitly test such mechanistic frameworks, our findings - particularly the increased expression of internally focused limbic–DMN states and reduced transition flexibility - may be interpreted as systems-level signatures of disrupted adaptive processing. These alterations could reflect a brain that is less responsive to exteroceptive demands and more bound by internally generated expectations or affective salience. Rather than implying that symptoms are “just psychological,” such an interpretation recognises the neurobiological basis of cognitive-emotional processing, the permeability of boundaries between organic and functional symptoms, and the importance of maintaining clinical sensitivity. We believe that adopting a cautious, integrative interpretive stance - grounded in systems neuroscience, informed by clinical experience, and open to multiple explanatory levels - provides the most constructive path forward for research in this area.

Exploratory correlation analyses provided partial support for the clinical relevance of altered brain-state dynamics in PCS. Specifically, the probability of occurrence of State 5 - a configuration dominated by limbic and default mode network (DMN) regions - was negatively associated with global cognitive performance on the MoCA and positively associated with serum levels of interleukin-1β (IL-1β), a key pro-inflammatory cytokine ^64, 65^. This state may reflect increased engagement of interoceptive and affectively laden neural circuits at the expense of externally directed cognitive functioning. Consistent with neuroimmune models of fatigue and sickness behaviour, sustained peripheral inflammation may bias intrinsic brain dynamics toward bodily monitoring and self-referential processing, potentially impairing cognitive flexibility and engagement with the external environment^66, 67^. However, the broader pattern of results warrants caution. More fine-grained analyses using the Cognitron platform, which assesses specific cognitive domains such as attention, working memory, and executive function, did not reveal significant associations with any LEiDA-derived dynamic metrics. Likewise, we found no robust relationships between dynamic state properties and self-reported symptoms, including fatigue and anxiety. These discrepancies suggest that while global measures such as the MoCA may capture diffuse effects of altered brain dynamics, these may not map neatly onto task-specific performance or subjective symptom ratings. It is possible that intrinsic brain dynamics are more closely linked to overall cognitive tone than to discrete cognitive operations or subjective complaints, or that limitations in power and measurement sensitivity may have obscured more subtle associations. Nonetheless, the observed link between State 5, inflammation, and global cognition supports the hypothesis that limbic-DMN dominance may represent a systems-level convergence point for neuroimmune signalling and cognitive disruption in PCS.

Our methodology offers both strengths and limitations. The use of multi-echo fMRI with ME-ICA denoising reduces susceptibility to motion and physiological noise, thereby improving signal quality for the analysis of dynamic functional connectivity. The LEiDA framework captures recurring whole-brain patterns of phase synchrony without the constraints of arbitrarily defined sliding windows, and allows for intuitive interpretation of transient brain states. Nonetheless, the sample size remains modest, and replication in larger cohorts is warranted. The resting-state acquisition duration (∼8 minutes) provides a reasonable estimate of baseline dynamics, although longer scans may increase sensitivity to rarer or transitional states. Group differences in educational attainment must also be acknowledged: although both groups fell within the high range of educational achievement, controls exhibited significantly higher levels on average. We chose not to include education as a covariate in primary analyses due to its collinearity with group status, which poses a risk of overcorrecting and obscuring meaningful group-related effects. Importantly, none of the brain dynamic metrics correlated significantly with years of education, suggesting that observed group differences are unlikely to be driven by this variable. Finally, while we anchored our state definitions to the Yeo 7-network atlas to facilitate interpretability, future studies employing finer-grained or individualized parcellations may help further refine the characterization of dynamic brain states.

In conclusion, our study reveals that PCS is characterised by a disruption in the brain’s intrinsic dynamic landscape - marked by diminished access to externally focused attention states and increased dwell in internally focused limbic-DMN configurations. This altered balance and reduced transition flexibility may underlie the persistent fatigue, attentional deficits, and emotional symptoms observed in PCS. Our results support a systems neuroscience account of PCS, implicating dynamic brain dysfunction in the sequelae of viral illness and identifying novel potential targets for monitoring and intervention.

## Credit authorship statement

**Marie-Stephanie Cahart:** Conceptualization, Methodology, Formal analysis, Writing – review & editing. **Owen O’Daly**: Methodology, Writing – review & editing. **Ziyuan Cai:** Data processing, Quality control, Writing – review & editing. **Nicole Mariani**: Methodological support, Writing – review & editing. **Alessandra Borsini**: Methodological support, Writing – review & editing. **Valeria Mondelli**: Conceptualization, Writing – review & editing. **Brandi Eiff**: Participant recruitment, Investigation, Data curation. **Silvia Rota**: Participant recruitment, Clinical assessments, Project coordination. **Timothy Nicholson**: Clinical oversight, Participant recruitment, Writing – review & editing. **Laila Raida**: Methodology, Writing – review & editing. **Adam Hampshire**: Cognitive task development, Normative modeling, Software, Writing – review & editing. **Fernando Zelaya**: Conceptualization, Methodology, Writing – review & editing. **Ottavia Dipasquale**: Methodology, Writing – review & editing. **Federico Turkheimer**: Statistical supervision, Interpretation, Writing – review & editing. **Steven C.R. Williams**: Resources, Funding acquisition, Supervision, Writing – review & editing. **Daniel Martins**: Conceptualization, Methodology, Formal analysis, Investigation, Writing – original draft, Writing – review & editing, Supervision, Project administration. All authors reviewed and approved the final manuscript and agree to be accountable for all aspects of the work.

## Acknowledgements

This study was funded by the National Institute for Health and Care Research (NIHR) Maudsley Biomedical Research Centre (BRC), South London and Maudsley NHS Foundation Trust, under the “Reach Out” funding call (Ref: R0-01). We are grateful to all participants for their time and commitment to the study. We thank the radiographers and technical staff at the Centre for Neuroimaging Sciences (King’s College London) for their assistance with MRI acquisition. We also acknowledge Synnovis (Viapath) for processing routine blood analyses, the Clinical Research Facility (CRF) team at King’s College Hospital Foundation Trust for their support during participant visits and blood collection, and the Cognitron platform team for assistance with cognitive data management and normative modelling.

## Conflict of interests

Nothing to declare.

## Notes

### Competing Interest Statement

The authors have declared no competing interest.

