## Supplementary for "Brain dynamics of attentional, default-mode and limbic networks are disrupted at rest in Post-COVID-19 Syndrome"

**Supplementary Table S1. Self-reported fatigue, affective symptoms, and somatic complaints in PCS and HC participants, with Bayesian inference.** Comparison of symptom severity between individuals with Post-COVID-19 Syndrome (PCS) and matched healthy controls (HC) across domains including fatigue (VAS, MFI), sleep (PSQI), autonomic function (COMPASS-31), post-exertional malaise, mood (HADS), PTSD, and fibromyalgia-related symptoms. Values are shown as mean (standard deviation), and group comparisons were conducted using independent samples t-tests or chi-squared tests, as appropriate. The Bayesian factor (BF₁₀) is reported for each comparison, quantifying the strength of evidence in favor of the alternative hypothesis. Asterisks (*) denote statistically significant differences at p < 0.05 or BF₁₀ > 1.

| **Variable** | **PCS**  **Mean (std)** | **HC**  **Mean (std)** | **Statistics** | **P-value** | **B_10_** |
| --- | --- | --- | --- | --- | --- |
| **Fatigue VAS** | 5.500 (1.606) | 1.850 (1.663) | T(38)=7.061 | 2.036x10^-8^* | 876.874* |
| **Multidimensional fatigue**  **General** **Physical**  **Reduced activity**  **Reduced motivation**  **Mental fatigue** | 15.200 (3.205)  14.600 (2.998)  12.150 (3.703)  8.600 (3.393)  12.850 (3.843) | 6.684 (2.212)  5.737 (2.884)  5.444 (2.307)  4.895 (2.378)  6.053 (3.027) | T(38)=47.68  T(38)=43.19  T(38)=15.09  T(38)=9.59  T(38)=21.41 | 1.355x10^-11*^  2.408x10^-11^*  1.072x10^-7*^  3.589x10^-4^*  4.391x10^-7^* | 1332.772*  4158.129*  650.746*  34.338*  767.860* |
| **ESS** | 12.400 (5.548) | 6.474 (4.234) | T(37)=3.735 | 6.304x10^-4^* | 24.874* |
| **HADS Depression** | 8.900 (2.954) | 4.053 (2.345) | T(37)=5.657 | 1.833x10^-6^* | 153.950* |
| **HADS Anxiety** | 7.550 (3.790) | 5.368 (2.477) | T(37)=2.116 | 0.041* | 1.287* |
| **PTSD total score** | 19.750 (10.208) | 6.105 (8.185) | T(37)=4.590 | 4.964x10^-5*^ | 59.715* |
| **De Paul's**  **PEM** | 2.950 (0.945) | 0.421 (0.449) | T(37)=10.584 | 9.541x10^-13*^ | 64546.865* |
| **De Paul's**  **Unrefreshing sleep** | 3.400 (0.821) | 1.316 (0.749) | T(37)=8.269 | 6.229x10^-10*^ | 1547.301* |
| **De Paul's**  **Autonomic symptoms** | 1.775 (0.881) | 0.553 (0.550) | T(37)=5.166 | 8.438x10^-6*^ | 73.625* |
| **De Paul's**  **Neurocognitive** | 1.775 (0.910) | 0.474 (0.612) | T(37)=5.212 | 7.316x10^-6*^ | 401.197* |
| **De Paul's**  **Immune** | 2.250 (0.993) | 0.658 (0.834) | T(37)=5.405 | 4.010x10^-6*^ | 381.905* |
| **COMPASS total score** | 62.413 (19.629) | 30.666 (19.158) | T(37)=5.108 | 1.011x10^-5*^ | 569.590* |
| **Global PSQI** | 15.200 (6.542) | 9.263 (5.791) | T(37)=2.995 | 0.005* | 15.254* |
| **MRC**  **Dyspnea**  **Scale** | Stage 1: 9  Stage 2: 9  Stage 3: 2 | 19  1  0 | χ^2^(5)=11.971 | 0.003* | N/A |
| **MoCA** | 26.8 (2.55) | 26.5 (1.54) | T(37)=0.430 | 0.668 | 0.420 |
| **Pain detect**  **WPI**  **SS** | 9.00(6.53)  5.45 (4.07)  8.55 (1.88) | 1.7 (1.84)  1.00 (1.62)  1.75 (1.68) | T(37)= 4.813  T(37)=4.541  T(37)=12.066 | <0.001*  <0.001*  <0.001* | 1315.956*  119.442*  13663.606* |
| **Fibromyalgia**  **diagnostic criteria** | Yes: 8  No: 12 | Yes:0  No:20 | χ^2^(2)=10.581 | 0.005* | N/A |

**Supplementary Table S2. Comparison of standard blood parameters between PCS and healthy control participants.** Descriptive statistics and group comparisons for hematological, biochemical, and inflammatory markers in individuals with Post-COVID-19 Syndrome (PCS) and matched healthy controls (HC). Results are reported as mean (standard deviation). Between-group comparisons were conducted using the Mann–Whitney U test (W), with corresponding p-values and Bayesian factors (BF₁₀) reported for each variable. Asterisks (*) denote statistically significant differences at p < 0.05 or BF₁₀ > 1.

| **Variable** | **PCS Mean (SD)** | **HC Mean (SD)** | **Statistics**  **W** | **P-value** | **B_10_** |
| --- | --- | --- | --- | --- | --- |
| **Sodium** | 140.118 (2.619) | 140.474 (2.091) | 161.00 | 1.00 | 0.369 |
| **Potassium** | 3.965 (0.237) | 4.000 (0.252) | 126.00 | 0.53 | 0.334 |
| **Urea** | 4.135 (1.346) | 4.337 (0.942) | 158.50 | 0.94 | 0.359 |
| **Creatinine** | 61.235 (8.318) | 68.632 (11.847) | 101.50 | 0.06 | 1.232* |
| **Glucose** | 4.871 (0.819) | 4.968 (1.106) | 157.00 | 0.90 | 0.329 |
| **Calcium_corrected** | 2.268 (0.059) | 2.288 (0.109) | 159.00 | 0.95 | 0.341 |
| **Phosphatase** | 1.088 (0.166) | 1.171 (0.163) | 106.00 | 0.08 | 0.749 |
| **Proteins_total** | 71.000 (3.691) | 71.526 (3.878) | 148.50 | 0.69 | 0.337 |
| **Albumin** | 47.000 (1.969) | 47.000 (2.749) | 154.50 | 0.83 | 0.326 |
| **Bilirubin_total** | 10.125 (6.672) | 9.824 (7.170) | 116.00 | 0.48 | 0.370 |
| **ALP** | 64.176 (15.204) | 63.947 (23.710) | 145.50 | 0.62 | 0.362 |
| **AST** | 23.353 (6.441) | 22.316 (7.521) | 131.00 | 0.34 | 0.420 |
| **Gamma_GT** | 13.059 (3.508) | 31.632 (47.978) | 97.50 | 0.04 | 1.735* |
| **Magnesium** | 0.859 (0.043) | 0.872 (0.054) | 151.50 | 0.76 | 0.347 |
| **CRP_recoded** | 1.353 (0.606) | 1.368 (0.597) | 158.50 | 0.92 | 0.397 |
| **Globulin** | 23.529 (3.875) | 24.526 (2.674) | 138.50 | 0.47 | 0.497 |
| **Ferritin** | 108.353 (103.511) | 99.789 (84.020) | 156.50 | 0.89 | 0.307 |
| **WCC** | 6.629 (1.662) | 6.501 (1.803) | 156.50 | 0.89 | 0.336 |
| **RCC** | 4.385 (0.457) | 4.561 (0.386) | 112.50 | 0.12 | 0.843 |
| **Hemoglobin** | 134.000 (13.224) | 137.263 (10.514) | 138.50 | 0.48 | 0.426 |
| **Haematocrit** | 0.403 (0.036) | 0.417 (0.029) | 120.00 | 0.19 | 0.739 |
| **MCV** | 91.988 (4.276) | 91.679 (5.684) | 159.00 | 0.95 | 0.329 |
| **MCH** | 30.624 (1.719) | 30.200 (2.149) | 151.00 | 0.75 | 0.348 |
| **MCHC** | 332.882 (11.931) | 329.158 (8.662) | 128.50 | 0.30 | 0.504 |
| **RDW** | 12.406 (0.599) | 12.474 (0.726) | 154.50 | 0.84 | 0.329 |
| **Platelet_count** | 267.118 (42.355) | 289.789 (45.281) | 105.00 | 0.08 | 1.123* |
| **MPV** | 10.988 (0.834) | 10.716 (0.727) | 121.00 | 0.20 | 0.463 |
| **Neutrophils** | 4.142 (1.178) | 3.773 (1.409) | 135.50 | 0.42 | 0.414 |
| **Lymphocytes** | 1.934 (0.550) | 2.029 (0.650) | 144.50 | 0.60 | 0.334 |
| **Monocytes** | 0.441 (0.102) | 0.441 (0.127) | 155.00 | 0.85 | 0.319 |
| **Eosinophils** | 0.141 (0.100) | 0.214 (0.243) | 147.50 | 0.67 | 0.365 |
| **Basophils** | 0.052 (0.030) | 0.042 (0.018) | 136.00 | 0.42 | 0.486 |
| **Immature_Granulocyte_Count** | 0.022 (0.016) | 0.022 (0.012) | 141.50 | 0.51 | 0.327 |
| **ESR** | 7.588 (6.662) | 8.278 (8.072) | 139.50 | 0.67 | 0.354 |
| **TSH** | 1.819 (0.630) | 1.633 (0.658) | 124.50 | 0.25 | 0.423 |
| **T4_free** | 15.124 (1.759) | 15.042 (1.624) | 156.00 | 0.87 | 0.321 |
| **NLR** | 2.209 (0.594) | 2.038 (1.137) | 125.50 | 0.26 | 0.540 |
| **MLR** | 0.234 (0.041) | 0.230 (0.072) | 156.00 | 0.87 | 0.346 |

**Supplementary Table S3. Serum cytokines and glial biomarkers in PCS and healthy control participants.** Group comparisons for circulating immune markers (interleukins, interferon-γ, TNF-α) and glial markers (S100β, GFAP) measured in serum samples from individuals with Post-COVID-19 Syndrome (PCS) and healthy controls (HC). Data are reported as mean (standard deviation). Concentrations are in pg/mL. Between-group differences were evaluated using the Mann–Whitney U test (W), with corresponding p-values and Bayes Factors (BF₁₀). Asterisks (*) indicate statistically suggestive effects (BF₁₀ > 1)

| **Variable** | **PCS Mean (SD)** | **HC Mean (SD)** | **Statistics**  **W** | **P-value** | **B_10_** |
| --- | --- | --- | --- | --- | --- |
| **IFNy** | 4.639 (8.032) | 4.182 (8.177) | 151.000 | 0.789 | 0.345 |
| **IL1b** | 0.345 (0.745) | 0.094 (0.061) | 195.000 | 0.300 | 0.573 |
| **IL6** | 1.197 (2.795) | 0.418 (0.205) | 194.000 | 0.478 | 0.431 |
| **TNFa** | 1.164 (1.799) | 0.980 (0.316) | 110.000 | 0.116 | 1.094* |
| **IL10** | 0.576 (1.145) | 0.229 (0.141) | 171.000 | 0.449 | 0.365 |
| **IL13** | 0.998 (0.889) | 1.228 (0.894) | 127.000 | 0.563 | 0.398 |
| **IL8** | 6.106 (4.671) | 6.581 (2.791) | 126.000 | 0.289 | 0.460 |
| **s100b** | 3.908 (2.192) | 5.165 (2.558) | 130.000 | 0.149 | 1.025* |
| **GFAP** | 14.62 (4.79) | 13.78 (5.32) | 157.000 | 0.515 | 0.406 |

**Supplementary Table S4. Performance on computerized cognitive tasks in PCS and healthy control participants.** Comparison of standardized deviation-from-expected (DFE) scores derived from the Cognitron cognitive task battery, assessing domains such as spatial reasoning, memory, motor control, and sustained attention. Scores represent z-standardized residuals relative to normative models controlling for age, sex, handedness, and ethnicity. Between-group differences were tested using independent samples t-tests, with corresponding p-values and Bayes Factors (BF₁₀). Asterisks (*) indicate results meeting the threshold for statistical significance (p < 0.05) or annedoctal Bayesian evidence (BF₁₀ > 1).

| **Variable** | **PCS Mean (SD)** | **HC Mean (SD)** | **Statistics** | **P-value** | **B_10_** |
| --- | --- | --- | --- | --- | --- |
| **Blocks_RT_DFE** | 0.136 (0.613) | -0.043 (0.539) | T(38)=0.981 | 0.33 | 0.452 |
| **BlocksSummaryScore_DFE** | -0.706 (1.069) | -0.301 (1.011) | T(38)=-1.231 | 0.23 | 0.563 |
| **Lead_Balloon_SummaryScore_DFE** | 0.576 (1.254) | -0.038 (0.658) | T(38)=1.939 | 0.06 | 1.330* |
| **2D_Manipulations_RT_DFE** | -0.006 (0.741) | 0.599 (1.892) | T(38)=-1.332 | 0.20 | 0.621 |
| **2D_Manipulations_SummaryScore_DFE** | 1.573 (1.484) | 1.236 (1.711) | T(38)=0.665 | 0.51 | 0.369 |
| **Motor_control_RT_DFE** | -0.097 (0.740) | -0.148 (0.766) | T(38)=0.214 | 0.83 | 0.314 |
| **Motor_Control_SummaryScore_DFE** | -0.012 (0.772) | -0.122 (0.764) | T(38)=0.453 | 0.65 | 0.335 |
| **Objects_memory_delayed_RT_DFE** | 0.416 (1.384) | -0.458 (0.898) | T(38)=2.369 | 0.02* | 2.653* |
| **Objects_memory_delayed_SummaryScore_DFE** | 0.073 (1.025) | -0.386 (1.866) | T(38)=0.964 | 0.34 | 0.446 |
| **Objects_memory_immediate_RT_DFE** | -0.106 (0.757) | -0.433 (0.840) | T(38)=1.293 | 0.20 | 0.596 |
| **Objects_memory_immediate_SummaryScore_DFE** | 0.023 (0.845) | -0.006 (1.268) | T(38)=0.085 | 0.93 | 0.310 |
| **Spotter_RT_DFE** | 0.360 (1.302) | -0.308 (0.841) | T(38)=1.927 | 0.06 | 1.303* |
| **Spotter_SummaryScore_DFE** | 0.093 (0.175) | 0.120 (0.361) | T(38)=-0.301 | 0.77 | 0.320 |
| **Verbal_analogies_RT_DFE** | 0.116 (0.743) | -0.027 (0.916) | T(38)=0.542 | 0.59 | 0.347 |
| **Verbal_analogies_SummaryScore_DFE** | -0.852 (1.355) | -0.592 (1.163) | T(38)=-0.651 | 0.52 | 0.366 |

**Supplementary Table S5. Correlations Between Brain States P1 and P5 and Clinical, Cognitive, and Biomarker Measures**. Partial Spearman correlations (Rho) between the fractional occupancy of dynamic brain states P1 and P5 and a range of clinical symptoms, cognitive performance scores, and peripheral biomarkers. All correlations were adjusted for group, age, gender, and dexterity. Corresponding p-values are reported for each Rho estimate.

| **Variable** | **Stats** | **P1** | **P5** |
| --- | --- | --- | --- |
| **MFI_General fatigue** | Rho | 0.211 | 0.212 |
|  | p-value | 0.225 | 0.223 |
| **MFI_Physical fatigue** | Rho | 0.283 | 0.029 |
|  | p-value | 0.099 | 0.869 |
| **MFI_Reduced activity** | Rho | 0.246 | 0.114 |
|  | p-value | 0.162 | 0.520 |
| **MFI_Reduced motivation** | Rho | 0.232 | 0.017 |
|  | p-value | 0.181 | 0.922 |
| **MFI_Mental fatigue** | Rho | -0.150 | 0.092 |
|  | p-value | 0.389 | 0.599 |
| **FAI_Total score** | Rho | -0.013 | 0.284 |
|  | p-value | 0.940 | 0.098 |
| **HADS_A_Total** | Rho | -0.009 | 0.106 |
|  | p-value | 0.960 | 0.545 |
| **HADS_D_Total** | Rho | 0.062 | 0.296 |
|  | p-value | 0.723 | 0.084 |
| **ESS_total** | Rho | -0.145 | 0.283 |
|  | p-value | 0.404 | 0.099 |
| **PTSD_Total** | Rho | -0.031 | 0.137 |
|  | p-value | 0.860 | 0.433 |
| **DP_PEM_Average_Severity** | Rho | -0.014 | 0.265 |
|  | p-value | 0.938 | 0.124 |
| **DP_Unrefreshing_Sleep_Average_Severity** | Rho | 0.339 | -0.030 |
|  | p-value | 0.047 | 0.864 |
| **DP_Autonomic_Average_Severity** | Rho | -0.304 | 0.113 |
|  | p-value | 0.076 | 0.519 |
| **DP_Neurocognitive_Average_Severity** | Rho | -0.189 | 0.291 |
|  | p-value | 0.276 | 0.090 |
| **DP_Immune_Average_Severity** | Rho | 0.063 | 0.192 |
|  | p-value | 0.720 | 0.270 |
| **Global_PSQI_Score** | Rho | -0.175 | 0.218 |
|  | p-value | 0.314 | 0.208 |
| **COMPASS_Total_Score** | Rho | 0.051 | 0.114 |
|  | p-value | 0.773 | 0.513 |
| **MOCA** | Rho | 0.127 | -0.400 |
|  | p-value | 0.474 | 0.019 |
| **PAIN detect score** | Rho | -0.248 | 0.116 |
|  | p-value | 0.144 | 0.502 |
| **WPI score** | Rho | 0.011 | 0.135 |
|  | p-value | 0.951 | 0.431 |
| **SS score** | Rho | 0.032 | 0.178 |
|  | p-value | 0.852 | 0.300 |
| **Time from 1st infection (months)** | Rho | 0.011 | 0.378 |
|  | p-value | 0.950 | 0.023 |
| **Fatigue VAS** | Rho | 0.247 | 0.018 |
|  | p-value | 0.147 | 0.916 |
| **IFNy** | Rho | 0.155 | 0.314 |
|  | p-value | 0.396 | 0.080 |
| **IL1b** | Rho | 0.074 | 0.363 |
|  | p-value | 0.688 | 0.041 |
| **IL6** | Rho | -0.069 | 0.122 |
|  | p-value | 0.704 | 0.500 |
| **TNFa** | Rho | 0.199 | 0.194 |
|  | p-value | 0.274 | 0.288 |
| **IL10** | Rho | 0.217 | -0.153 |
|  | p-value | 0.241 | 0.412 |
| **IL13** | Rho | 0.092 | -0.210 |
|  | p-value | 0.629 | 0.266 |
| **IL8** | Rho | -0.104 | -0.174 |
|  | p-value | 0.570 | 0.342 |
| **s100b** | Rho | 0.090 | 0.171 |
|  | p-value | 0.614 | 0.335 |
| **GFAP** | Rho | -0.050 | -0.088 |
|  | p-value | 0.779 | 0.622 |
| **Sodium** | Rho | -0.020 | 0.015 |
|  | p-value | 0.913 | 0.933 |
| **Potassium** | Rho | -0.196 | 0.042 |
|  | p-value | 0.299 | 0.825 |
| **Urea** | Rho | -0.031 | 0.123 |
|  | p-value | 0.868 | 0.502 |
| **Creatinine** | Rho | 0.130 | 0.143 |
|  | p-value | 0.477 | 0.435 |
| **Glucose** | Rho | -0.160 | 0.026 |
|  | p-value | 0.382 | 0.889 |
| **Calcium_corrected** | Rho | 0.029 | -0.108 |
|  | p-value | 0.877 | 0.558 |
| **Phosphatase** | Rho | -0.124 | 0.089 |
|  | p-value | 0.500 | 0.628 |
| **Proteins_total** | Rho | -0.119 | -0.368 |
|  | p-value | 0.516 | 0.038 |
| **Albumin** | Rho | 0.142 | -0.160 |
|  | p-value | 0.438 | 0.380 |
| **Bilirubin_total** | Rho | -0.396 | -0.008 |
|  | p-value | 0.033 | 0.966 |
| **ALP** | Rho | 0.026 | 0.007 |
|  | p-value | 0.886 | 0.972 |
| **AST** | Rho | -0.012 | -0.026 |
|  | p-value | 0.948 | 0.889 |
| **Gamma_GT** | Rho | -0.235 | 0.079 |
|  | p-value | 0.195 | 0.667 |
| **Magnesium** | Rho | -0.180 | 0.132 |
|  | p-value | 0.324 | 0.471 |
| **Globulin** | Rho | -0.154 | -0.219 |
|  | p-value | 0.399 | 0.229 |
| **Ferritin** | Rho | 0.039 | 0.081 |
|  | p-value | 0.834 | 0.660 |
| **WCC** | Rho | 0.226 | -0.111 |
|  | p-value | 0.214 | 0.545 |
| **RCC** | Rho | 0.155 | -0.013 |
|  | p-value | 0.397 | 0.942 |
| **Hemoglobin** | Rho | 0.150 | 0.050 |
|  | p-value | 0.413 | 0.784 |
| **Haematocrit** | Rho | 0.138 | 0.050 |
|  | p-value | 0.452 | 0.785 |
| **MCV** | Rho | -0.121 | 0.131 |
|  | p-value | 0.510 | 0.476 |
| **MCH** | Rho | 0.054 | 0.002 |
|  | p-value | 0.767 | 0.990 |
| **MCHC** | Rho | 0.115 | -0.122 |
|  | p-value | 0.532 | 0.505 |
| **RDW** | Rho | -0.169 | 0.040 |
|  | p-value | 0.355 | 0.827 |
| **Platelet_count** | Rho | -0.094 | 0.068 |
|  | p-value | 0.607 | 0.710 |
| **MPV** | Rho | 0.093 | -0.180 |
|  | p-value | 0.612 | 0.324 |
| **Neutrophils** | Rho | 0.100 | -0.149 |
|  | p-value | 0.587 | 0.415 |
| **Lymphocytes** | Rho | 0.291 | -0.130 |
|  | p-value | 0.106 | 0.478 |
| **Monocytes** | Rho | 0.237 | 0.043 |
|  | p-value | 0.191 | 0.816 |
| **Eosinophils** | Rho | 0.035 | -0.018 |
|  | p-value | 0.847 | 0.921 |
| **Basophils** | Rho | 0.078 | -0.230 |
|  | p-value | 0.671 | 0.205 |
| **Immature_Granulocyte_Count** | Rho | -0.080 | -0.159 |
|  | p-value | 0.665 | 0.384 |
| **ESR** | Rho | -0.152 | -0.241 |
|  | p-value | 0.414 | 0.191 |
| **TSH** | Rho | -0.186 | -0.002 |
|  | p-value | 0.307 | 0.993 |
| **T4_free** | Rho | -0.015 | 0.086 |
|  | p-value | 0.933 | 0.639 |
| **NLR** | Rho | -0.166 | -0.028 |
|  | p-value | 0.364 | 0.878 |
| **MLR** | Rho | -0.110 | 0.302 |
|  | p-value | 0.550 | 0.093 |
| **Blocks_RT_DFE** | Rho | 0.128 | -0.139 |
|  | p-value | 0.458 | 0.420 |
| **BlocksSummaryScore_DFE** | Rho | 0.106 | -0.205 |
|  | p-value | 0.540 | 0.229 |
| **Lead_Balloon_SummaryScore_DFE** | Rho | 0.125 | -0.137 |
|  | p-value | 0.467 | 0.425 |
| **2D_Manipulations_RT_DFE** | Rho | 0.167 | 0.042 |
|  | p-value | 0.331 | 0.806 |
| **2D_Manipulations_SummaryScore_DFE** | Rho | -0.172 | 0.024 |
|  | p-value | 0.315 | 0.888 |
| **Motor_control_RT_DFE** | Rho | 0.232 | -0.220 |
|  | p-value | 0.173 | 0.197 |
| **Motor_Control_SummaryScore_DFE** | Rho | 0.234 | -0.192 |
|  | p-value | 0.169 | 0.262 |
| **Objects_memory_delayed_RT_DFE** | Rho | 0.005 | 0.038 |
|  | p-value | 0.977 | 0.828 |
| **Objects_memory_delayed_SummaryScore_DFE** | Rho | 0.057 | -0.107 |
|  | p-value | 0.741 | 0.533 |
| **Objects_memory_immediate_RT_DFE** | Rho | 0.138 | -0.181 |
|  | p-value | 0.423 | 0.292 |
| **Objects_memory_immediate_SummaryScore_DFE** | Rho | -0.074 | -0.146 |
|  | p-value | 0.667 | 0.396 |
| **Spotter_RT_DFE** | Rho | 0.005 | 0.034 |
|  | p-value | 0.978 | 0.842 |
| **Spotter_SummaryScore_DFE** | Rho | -0.080 | 0.136 |
|  | p-value | 0.643 | 0.428 |
| **Verbal_analogies_RT_DFE** | Rho | -0.013 | -0.107 |
|  | p-value | 0.940 | 0.536 |
| **Verbal_analogies_SummaryScore_DFE** | Rho | -0.002 | 0.168 |
|  | p-value | 0.989 | 0.327 |
